# Identification and targeting of a novel drug pocket in the tumour angiogenesis regulator RhoJ

**DOI:** 10.64898/2026.10.14.758331

**Authors:** Wenxuan Chen, Deborah K. Shoemark, Robin A. Corey, Tatyana N. Andrienko, Harry Mellor

**Affiliations:** School of Biochemistry and Biomedical Sciences, University of Bristol, University Walk, Bristol BS8 1TD; School of Psychology and Neuroscience, University of Bristol, University Walk, Bristol BS8 1TD

**Keywords:** vascular pharmacology, drug discovery, tumour angiogenesis, Rho GTP-binding proteins, molecular dynamics simulation, bioluminescence resonance energy transfer

## Abstract

The Rho GTPase RhoJ is a key regulator of tumour angiogenesis and has been identified as a target for anti-angiogenic drug design. Its high sequence conservation with the related Rho GTPase Cdc42 complicates selective drug discovery. RhoJ has a unique N-terminal extension (Asn2-Lys19) that is absent in Cdc42 and other Rho GTPases. Here, we use molecular dynamics simulations to identify a novel allosteric pocket formed by the RhoJ N-terminus and show that this pocket contributes to interactions with key downstream signalling partners. Structure-based virtual screening of small molecules targeting this pocket identified the antifungal terconazole as a potential binder. Computational docking revealed that terconazole occupies the N-terminal pocket, sharing critical contacts with the interface between RhoJ and PAK3, a key interacting partner. To validate this, we developed an optimized bioluminescence resonance energy transfer (eBRET^2^)-based biosensor to monitor the interaction between RhoJ and PAK3 in live cells and saw that terconazole disrupted the RhoJ:PAK3 interaction in a dose-dependent manner. Together, our MD results show that the RhoJ N-terminal extension assumes structure on binding to PAK3 and contributes to that interaction, while our cellular biosensor data support targeting this pocket to modulate the RhoJ:PAK3 interaction. The ability of terconazole to bind to the N-terminal pocket and disrupt PAK3 binding provides proof of concept for the specific targeting of RhoJ in the development of novel anti-angiogenic drugs.

**Graphical Abstract:** 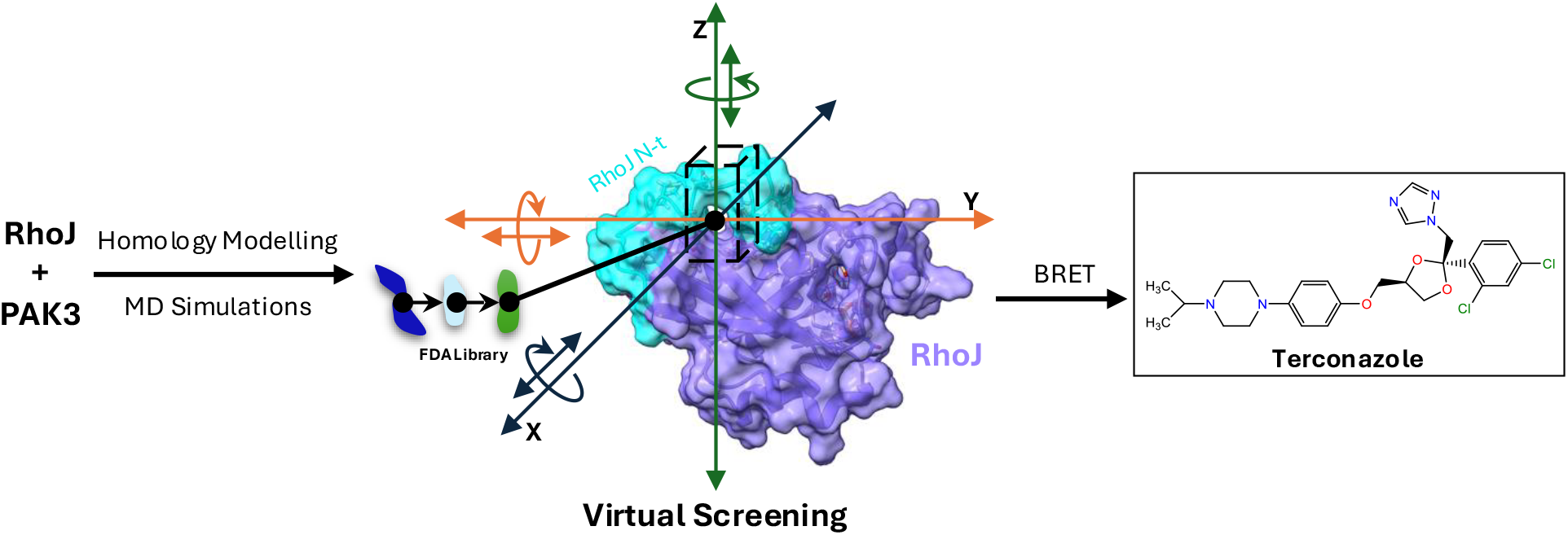

## 1. Introduction

The Rho GTPase family are the major regulators of cell shape and cell migration in eukaryotic cells [1]. The human Rho GTPase family comprises 20 members, of which RhoA, Rac1 and Cdc42 are the best characterized [2]. Humans have two Rho GTPases that are closely related to Cdc42, and which arose by duplication of the Cdc42 gene during chordate evolution – RhoQ and RhoJ [3]. Whereas Cdc42 is expressed globally in human tissues, the distributions of RhoQ and RhoJ are more restricted. RhoQ has roles in neuronal morphology and axonal outgrowth [4]. RhoJ is expressed in endothelial cells under control of the ERG transcription factor [5]. Mice lacking RhoJ are viable but have defects in blood vessel formation during development [6]. Specifically, RhoJ is required for angiogenesis [7, 8], the process by which new vessels sprout from preexisting vessels. This process is highly active during development and is responsible for the elaboration of most of the vasculature [9, 10]. In adult humans, angiogenesis is mostly dormant, but can be reinitiated in response to wound healing, and tissue hypoxia.

Angiogenesis is also activated in cancer, in the early stages of tumour formation [9, 10]. As the tumour grows, cancer cells at the core become hypoxic, initiating the natural angiogenic response. Cancer cells also produce pro-angiogenic factors to further stimulate blood vessel formation. This tumour angiogenesis provides the growing mass with oxygen and nutrients, supporting further expansion. Tumour vessels also provide routes for metastatic cells to enter the circulatory system and to travel to distant sites, seeding secondary tumour growth. Drugs that inhibit tumour angiogenesis, such as Bevacizumab, are now widely used in cancer treatment, driving regression of tumour vessels and shrinkage of the tumour mass [11].

Bevacizumab and related drugs are antibody-based therapeutics that work by sequestering pro-angiogenic factors secreted by the tumour cells [12]. They have been successful in clinical use; however, they have some features that limit their effectiveness [11]. As with all antibody-based therapeutics, they are poorly accessible to dense tissue such as tumours.

Responsiveness is limited to tumour angiogenesis driven by the target of these drugs – vascular endothelial growth factor (VEGF). Further, resistance develops when tumour cells begin to secrete other pro-angiogenic factors [13]. Recognition of the limitations of current anti-angiogenic drugs has led to the search for additional targets and approaches in the control of tumour angiogenesis [14].

RhoJ signals downstream of VEGF but also mediates angiogenesis in response to other pro-angiogenic factors [15]. The central role of RhoJ in pro-angiogenic signalling [5, 7, 16-20], and the seeming restriction of RhoJ to endothelial cells [5], make it an attractive target for the development of novel inhibitors of tumour angiogenesis. RhoJ has been shown to be required for tumour angiogenesis in mice, and genetic targeting of RhoJ leads to disruption of pre-existing peritumoural vessels and inhibition of tumour angiogenesis [8]. Historic attempts to generate inhibitors of Rho GTPases have met with mixed results. Initial work focused on RhoA, Rac1 and Cdc42, due to their key roles in cancer cell invasion and metastasis. Similar to attempts to target the Ras GTPase, it proved difficult to identify suitable pockets on the small GTPase surface that could be targeted by small molecules. The first successful compounds identified worked by targeting upstream activators of those Rho GTPases [21].

Screening for small molecules that could compete for nucleotide binding on Cdc42 identified ML141, which functionally inhibits GTP binding to Cdc42 through a noncompetitive/ allosteric mechanism [22]. Subsequent work has identified additional direct inhibitors of Cdc42, several of which show apparent specificity over other Rho GTPases [21]. Recent work by Jahid *et al*. identified a novel pocket on the surface of Cdc42 and then used structure-based virtual screening (SBVS) to identify small molecule inhibitors that targeted that pocket [23]. The pocket was shown to also be present in RhoJ. These compounds have the potential to be used as RhoJ inhibitors in tumour angiogenesis; however, the global expression of Cdc42 and its central role in a multitude of critical cellular processes mean that the toxicity caused by such dual-targeting drugs would be prohibitive [24]. There is a clear rationale for the development of specific small molecule inhibitors of RhoJ, and this requires approaches that target features of this signalling protein that are absent from Cdc42.

We used atomistic molecular dynamics (MD) simulations to identify a novel pocket in RhoJ formed by its unique N-terminal extension when bound to its downstream signalling partner PAK3. We performed a virtual screen to identify potential hit compounds that bind to this site and developed a cell-based BRET assay to test the ability of those compounds to disrupt the RhoJ:PAK3 interaction. Our data reveal that the antifungal compound terconazole can bind to the RhoJ pocket and disrupt RhoJ binding to PAK3, providing proof of concept for future therapeutic development.

## 2. Materials and Methods

### Multiple sequence alignment

Multiple sequence alignments of Cdc42 subfamily GTPases [Cdc42 (UniProt: P60953), RhoQ (UniProt: P17081), RhoJ (UniProt: Q9H4E5)] were performed using T-Coffee [25] with default parameters.

### In silico protein structure preparation and homology modelling

Homology models for RhoJ-Mg^2+^-GppCp in complex with p21-binding domains (PBDs) from 4 effector proteins [PAK1 (UniProt: Q13153; a.a. 75-118), PAK3 (UniProt: O75914-2; a.a. 65-108), N-WASP (UniProt: O00401; a.a. 195-253), and BORG3 (UniProt: Q6NZY7; a.a. 14-62)] were generated using UCSF Chimera(version 1.17.1, University of California, San Francisco, CA, USA; [26]). The Cdc42-Mg^2+^-GppNHp:PBD^PAK1^ structure (PDB: 1E0A) served as the template for modelling the RhoJ-Mg^2+^-GppCp:PBD^PAK1^ complex, the RhoJ-Mg^2+^-GppCp:PBD^PAK3^ complex, and the Cdc42-Mg^2+^-GppNHp:PBD^BORG3^ complex. The Cdc42-Mg^2+^-GppCp:PBD^WASP^ structure (PDB: 1CEE) served as the template for modelling the Cdc42-Mg^2+^-GppCp:PBD^N-WASP^ complex and the RhoJ-Mg^2+^-GppCp:PBD^N-WASP^ complex. The Cdc42-Mg^2+^-GppCp:PBD^PAK6^ structure (PDB: 2ODB) served as the template for RhoJ-Mg^2+^-GppCp:PBD^BORG3^ complex. GppCp and GppNHp were parameterized using ACPYPE [27]. Models were subjected to structural alignment in Chimera, and hydrogen atoms were added using pdb2gmx in the GROMACS version 2022.6 suite of software [28], with standard protonation states assigned for ionisable side chains at physiological pH. Histidine protonation states were assigned by pdb2gmx based on the local hydrogen-bonding environment.

Membrane-associated systems were constructed using CHARMM-GUI [29]. A farnesyl group was attached to the C-terminal Cys211 of RhoJ and embedded into a cholesterol:POPC:POPE (1:4:4) bilayer, with or without PBD^PAK3^. To further examine the N-terminal behaviour of RhoJ in a lipidated context, additional models were generated with N-terminal palmitoylation at Cys3 and Cys11. Palmitate, and farnesyl were also parameterized using ACPYPE [27].

### Molecular dynamics (MD) simulations

All simulations were set up and performed using GROMACS (version 2022.6), with soluble protein models parameterized using the AMBER99SB-ILDN force field and membrane-associated models using AMBER99SB-ILDN_SLIPIDS forcefield. Membrane-free systems were solvated in explicit TIP3P water and neutralized with 150 mM NaCl. Systems were energy-minimized in a triclinic box using the steepest descent algorithm (2000 steps; maximum force tolerance of 200 kJ•mol^-1^•nm^-1^), followed by validations via PROCHECK to ensure stereochemical quality. Following 50 ps of NPT position-restrained equilibration, three independent 500 ns production runs were performed for each complex at 310 K (velocity-rescale method) and 1 bar (Berendsen barostat). Post-MD analysis and residue interaction inspections were conducted using GROMACS tools, LigPlot+ [30] (version 2.2), and ChimeraX [31] (version 1.8, University of California, San Francisco, CA, USA). Predicted solvation free-energy gains upon interface formation (Δ^i^G) were calculated using PDBePISA [32].

### Computational alanine scanning (CAS) and virtual screening

Hot-spot residues in the complexes of interest were identified via computational alanine scanning using the Bristol University Docking Engine (BUDE) Alanine Scan (BAlaS) web server [33]. This approach systematically calculates the change in binding free energy (ΔΔG) resulting from the individual mutation of interfacial residues to alanine. Residues with more positive ΔΔG values are predicted to make greater contributions to the stability and binding affinity of the target complex.

Virtual screening was performed using BUDE [34]. A library of 1,537 FDA-approved drugs (e-Drug3D [35]; version 2022.07) was docked into the RhoJ N-terminal pocket using an ensemble of 5 representative frames sampled across the stable MD trajectory window (200– 500 ns) of the RhoJ-Mg^2+^-GppCp:PBD^PAK3^ MD replicate that showed the lowest overall RMSD_Cα_ across the 500-ns simulation. For each frame, docking was performed within a 3 × 3 × 9 Å grid centred on PAK3 Leu72–Pro73–Ser74 in the RhoJ:PBD^PAK3^ complex using the BUDE evolutionary Monte Carlo algorithm, allowing 3 translational (X, Y, Z) and 3 rotational (X_Rot_, Y_Rot_, Z_Rot_) degrees of freedom. BUDE-predicted binding energies were averaged across the 5 frames to rank candidates.

### Cell culture and transfection

HEK293T human embryonic kidney cells (American Type Culture Collection) were cultured in high glucose Dulbecco′s modified Eagle′s medium (DMEM-high glucose; D5796-500ML, Sigma-Aldrich, St. Louis, MO, USA) supplemented with 10% foetal bovine serum (FBS; F9665, Sigma-Aldrich), 50 U/mL penicillin and 50μg/mL streptomycin (15070063, Thermo Fisher Scientific, Waltham, MA, USA), at 37°C in a humidified atmosphere of 5% CO_2_. To express RhoJ biosensor, donor (RhoJ_Q79L_-RLuc8-Ct) and acceptor (PBD^PAK3^-GFP^2^ or GFP^2^-PBD^PAK3^) cDNAs were cloned into pLVX-Puro (Clontech Laboratories, Inc., Mountain View, CA, USA). Cells were transfected with plasmids of interest (at appropriate donor:acceptor ratios required for the purpose of the assay) at 60% confluence using Lipofectamine™ 2000 (11668-019, Invitrogen, Carlsbad, CA, USA) according to the manufacturer’s instructions. Cells were used for experiments 48 h post-transfection.

### Bioluminescence Resonance Energy Transfer assays

HEK293T cells expressing the eBRET^2^ biosensor or a donor-only control were seeded at a density of 100,000 cells/well in white 96-well Nunclon™ Delta-treated flat-bottom plates (136101, Thermo Scientific) at a donor-to-acceptor ratio of 1:10 (or appropriate ratios for transfection optimization of the biosensor and donor saturation curve determination). Cells were cultured for 24 h in BRET assay medium [HEPES-buffered DMEM-HG (21063-029, Gibco, Grand Island, NY, USA) supplemented with 2 mM L-glutamine (25030-024, Gibco™), 50 U/mL penicillin, 50 μg/mL streptomycin] supplemented with 10% (v/v) FBS at 37°C in a 5% CO_2_ atmosphere. One hour prior to stimulation, the culture medium was replaced with fresh BRET assay medium. Cells were subsequently treated with the indicated test compounds [DMSO (2650-5 × 5ML, Sigma-Aldrich); ML141 (SML0407-5MG, Sigma Aldrich); Cilostazol (PHR1503-1G, Sigma-Aldrich); Capmatinib (HY-13404-10mg, MedChemexpress LLC, Monmouth Junction, NJ, USA); Tucatinib (7640/10, Tocris Bioscience™, Bio-techne, Bristol, UK); Avapritinib (BLU-285) (22985-1 mg, Cayman Chemical, Ann Arbor, MI, USA); Losartan potassium salt (462740050, Thermo Scientific); Losartan carboxylic acid (SMB01354-5MG, Sigma-Aldrich); Terconazole (32355-100MG, Sigma-Aldrich)] and incubated at 37°C as required. BRET measurements were performed using a Synergy Neo2 hybrid multimode plate reader (Agilent Technologies, Santa Clara, CA, USA). The system automatically injected the coelenterazine 400a substrate (5 μM final concentration) into each well. Luminescence was recorded at 410/80 nm (donor emission) and 515/30 nm (acceptor emission) at 16.30 s post-injection. All experiments were conducted in triplicate and repeated in 3 independent biological replicates.

The raw BRET ratio was defined as the ratio of the acceptor emission (I_acceptor_) to the donor emission (I_donor_): Raw BRET Ratio (BU) =I_acceptor_/I_donor_. To account for background interference, the corrected BRET ratio (BRET_corrected_) was calculated by subtracting the mean BRET ratio of cells expressing only the donor from the raw BRET ratio of the biosensor pair: 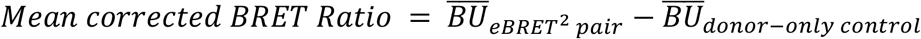. Donor saturation curve, dose-response curves and IC_50_ values were determined using nonlinear regression or four-parameter logistic (4PL) regression analysis in GraphPad Prism 10.1.2.

### Cdc42 inhibition assays

Inhibition of Cdc42 binding to PAK1 was measured using the G-LISA® Cdc42 Activation Assay Biochem Kit™ (BK127, Cytoskeleton Inc., Denver, CO, USA). 2.4 ng of purified recombinant constitutively active Cdc42 protein (CDCA, Cytoskeleton Inc.) in lysis buffer (GL35, Cytoskeleton Inc.) was treated with 5% (v/v) DMSO or terconazole for 15 min at 4°C. Following treatment, Cdc42 binding to the immobilised PBD^PAK1^ was quantified according to the manufacturer’s instructions. Absorbance was measured at 490 nm using a POLARstar Omega microplate spectrophotometer (BMG LABTECH, Ortenberg, Germany).

### Quantification and statistical analysis

GraphPad Prism 10.1.2 (GraphPad Software, Boston, MA, USA) was used to perform all quantification and statistical analysis of data. Data are presented as means ± SEM. Significant differences between two groups were determined using unpaired two-tailed Welch’s t-tests.

Experiments with a single independent variable with more than two sample groups use one-way ANOVA to compare the variance in the group means within a sample, followed by Bonferroni’s or Dunnett’s post-hoc test, as indicated. Experiments with at least two independent variables use two-way ANOVA to examine the effect of 2 factors using the Bonferroni test, as indicated. * *P* < 0.05; ** *P* < 0.01, ** *P* < 0.001; *** *P* < 0.001; **** *P* < 0.0001.

## 3. Results

### 3.1 The RhoJ N-terminal extension forms a pocket upon PAK3 binding

RhoJ and RhoQ are highly similar to Cdc42 in their core globular domain, with sequence identities of 63% and 65%, respectively. Sequence alignment shows that the core G-boxes (G1–G5) and switch regions remain highly conserved, with variation mostly confined to the C-terminal hypervariable region (Fig. 1A). Additionally, RhoJ possesses a distinct N-terminal region (Met1–Lys20), within which residues 2–19 form a RhoJ-specific N-terminal extension relative to Cdc42 and RhoQ (Fig. 1A). The function of this region is unclear, but it has previously been shown to affect the membrane localization and nucleotide loading of RhoJ [36].

**Fig. 1.**
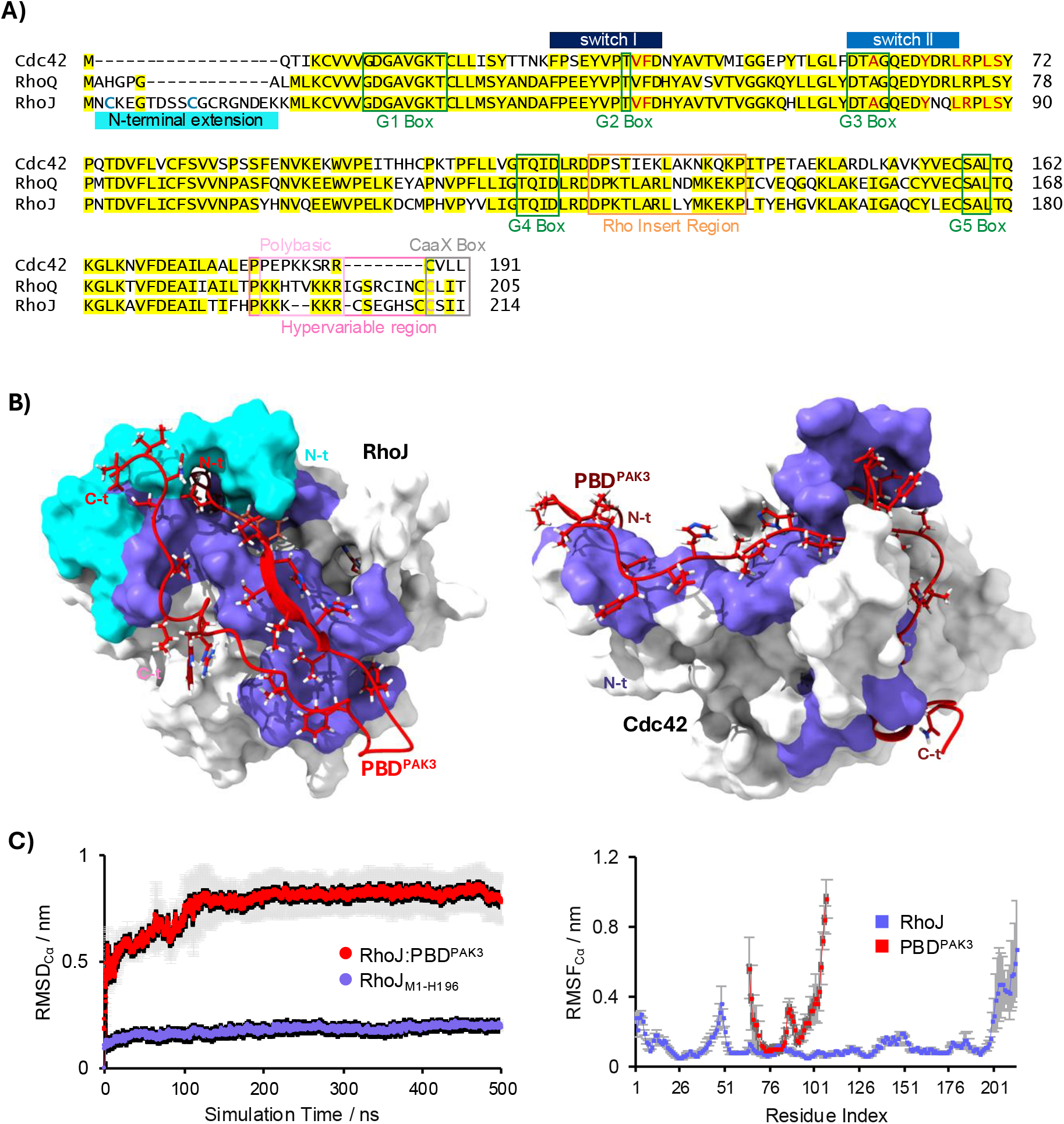
Allosteric N-terminal pocket of RhoJ revealed by MD modeling. **(A)** Multiple-sequence alignment (MSA) of human Cdc42 subfamily GTPases (Cdc42, RhoQ, and RhoJ) performed with T-Coffee. Residues are shaded by conservation (yellow). Structural and regulatory motifs are indicated: G-boxes G1–G5 (green), Switch Region I (dark blue), Switch Region II (blue), Rho-insert region (tan), hypervariable C-terminus (hot pink), polybasic region (light pink), and CaaX box (grey). The RhoJ-specific N-terminal extension is highlighted (cyan). Cysteine residues are colored by predicted post-translational modification: S-palmitoylation (dark cyan), S-farnesylation (rose), and S-geranylgeranylation (olive). Residues comprising the previously predicted allosteric binding pocket are colored in dark red. Representative structures of activated GTPases complexed with the PBD^PAK3^. (Left) Lowest-RMSD_Cα_ replica of the RhoJ-Mg^2+^-GppCp:PBD^PAK3^ complex from all-atom MD simulations at 500 ns. (Right) NMR structure of the Cdc42:PBD^PAK3^ complex (PDB: 1EES) for comparison. GTPases are shown in surface representation (core: white; RhoJ N-terminus Met1-Lys20: cyan; PBD^PAK3^ interface: medium slate blue). PBD^PAK3^ is shown as a red ribbon/stick model. **(B)** MD stability and flexibility of the activated RhoJ:PBD^PAK3^ complex. Left, mean RMSD_Cα_ over 0-500 ns; right, per-residue RMSF_Cα_ over the equilibrated window (150-500 ns). Data are mean ± SEM (*n* = 3 independent simulations).

In our previous studies, we showed that Cdc42 and RhoJ bind to an overlapping set of protein partners, but with seemingly different affinities [16]. We were interested in the structural basis of these differences, as they control the specificity of signalling. There are currently no available structures of RhoJ, alone or in complex with its binding partners. To address this, we constructed a homology model of the activated RhoJ-Mg^2+^-GppCp:PBD^PAK3^ complex. We found previously that PAK3 was the predominant signalling partner for RhoJ in endothelial cells [16]. Due to the absence of ligand information in the Cdc42:PAK3 NMR structure (PDB: 1EES), we used the Cdc42-Mg^2+^-GppNHp:PBD^PAK1^ NMR structure (PDB: 1E0A) as the structural template, given the high sequence homology between RhoJ and Cdc42 in the switch regions and the structural similarity between PBD^PAK3^ and PBD^PAK1^. The RhoJ sequence, including its unique N-terminal extension, was aligned onto the Cdc42 template, and the resulting homology model was energy-minimised prior to simulation. All-atom MD simulations of the RhoJ-Mg^2+^-GppCP:PBD^PAK3^ complex were subsequently performed to assess the conformational behaviour of the N-terminal extension under physiological simulation conditions.

Analysis of the MD trajectories across 3 independent simulation replicates revealed that the N-terminal region (cyan) does not remain disordered in the context of the complex; instead, it consistently converges into a defined conformation across all 3 replicates, facilitating the reproducible formation of a novel binding pocket at the PBD^PAK3^ interface (Fig. 1B; left). To contextualise this observation, the equilibrated RhoJ complex was aligned against the Cdc42:PBD^PAK3^ NMR structure (PDB: 1EES) (Fig. 1B; right). This comparison demonstrated a markedly different surface topology at the equivalent effector interface in Cdc42 and confirmed that the pocket is a structural feature of PAK3 binding that is unique to RhoJ (Fig. 1B). The stability of the simulated RhoJ-Mg^2+^-GppCp:PBD^PAK3^ complex was evaluated by monitoring the RMSD_Cα_ over the full 500 ns trajectory (Fig. 1C). Following an initial equilibration phase, the overall complex reached a stable plateau at approximately 0.8 nm, whereas the RhoJ core segment containing its N-terminal extension (Met1–His196) remained highly rigid (RMSD_Cα_ < 0.2 nm), indicating the robust conformational stability of the folded N-terminal pocket (Fig. 1C). Per-residue RMSF_Cα_ also demonstrated the structural persistence of the pocket region, with a low mobility and low fluctuation values observed across both the RhoJ core and its N-terminus throughout the simulation (Fig. 1C).

Detailed structural characterization of this interface identified a cluster of interaction hotspots. Specifically, RhoJ N-terminal residues form stable hydrogen bonds and hydrophobic contacts with PAK3 residues Leu72, Pro73, and Ser74 (Fig. 2A, B). To inspect the energetic contribution of these residues, we performed computational alanine scanning (CAS). Comparison of per-residue binding energies between RhoJ:PBD^PAK3^ and Cdc42:PBD^PAK3^ further corroborated the energetic contribution of PAK3 Leu72-Ser74 to the formation of the RhoJ N-terminal pocket, which was also manifested in pronounced per-residue displacements (Fig. 2C). From the RhoJ perspective, the RhoJ N-terminus (M1-K20) contributed ∼19.2% binding energy to the RhoJ:PBD^PAK3^ interaction (Fig. 2D). Direct *in silico* truncation of the RhoJ N-terminal region (M1-K20) from MD-derived RhoJ:PBD^PAK3^ models demonstrated that RhoJ N-terminus accounts for 285.6 Å^2^ binding interface, corresponding to 17.0% of the total RhoJ:PBD^PAK3^ interface (Fig. 2E). To evaluate this interface under biomimetic conditions, membrane-associated RhoJ:PBD^PAK3^ models were constructed using CHARMM-GUI, incorporating dual N-terminal palmitoylation (Cys3 and Cys11) [36] and C-terminal farnesylation (Cys211) [37]. In these membrane-embedded simulations, the N-terminal region consistently maintained a substantial interaction footprint, contributing 213.4 Å^2^, representing 14.0% of the total interface area (Fig. 2F). Furthermore, N-terminal S-palmitoylation of RhoJ reduced the magnitude of the predicted solvation free-energy gain upon interface formation (|Δ^i^G|) by approximately 17.7% (Fig. 2G) calculated by PDBePISA [32], consistent with the substantial energetic contribution of the RhoJ N-terminus to this interaction identified independently by BUDE computational alanine scanning (Fig. 2D). Collectively, these thermodynamic and spatial evaluations demonstrate that the RhoJ N-terminus provides a highly reproducible, structurally stable contribution to the PAK3-binding interface across soluble and membrane-associated models. All of these suggest the presence of a stable, druggable allosteric site unique to RhoJ, which is suitable for targeted inhibition.

**Figure 2.**
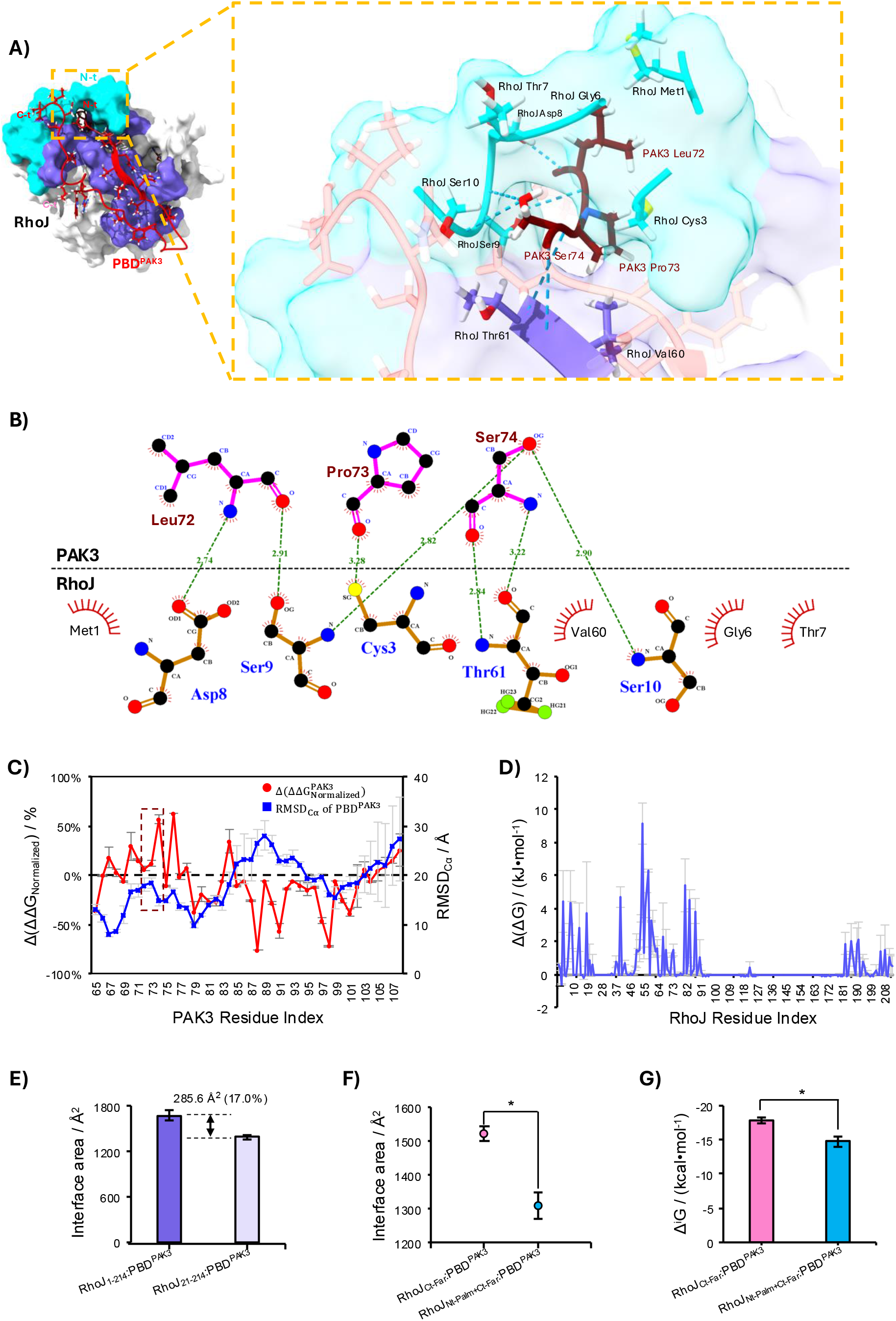
Structural and energetic characterization of the RhoJ N-terminal pocket interface. **(A)** Close-up 3D view of interaction hotspots within the RhoJ:PBD^PAK3^ N-terminal pocket from the 500 ns MD simulation frame. PAK3 residues Leu72, Pro73, and Ser74 are highlighted in maroon; the RhoJ N-terminus is cyan, and interacting residues are medium slate blue. Cyan dashed lines indicate hydrogen bonds. **(B)** 2D interaction map of N-terminal pocket hotspots. The black dashed line represents the protein-protein interface. Hydrogen bonds are shown as green dotted lines (lengths in Å); hydrophobic interactions are indicated by red radial sectors. **(C)** Comparative computational alanine scanning (CAS) of PBD^PAK3^ binding to RhoJ and Cdc42 with quantification of displacement of PBD^PAK3^. The red line represents 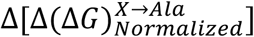, calculated as the difference in average normalized per-residue binding energy change 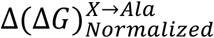 between the RhoJ:PBD^PAK3^ MD models and the Cdc42:PBD^PAK3^ NMR structure (PDB: 1EES) 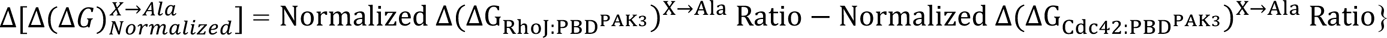. Blue points indicate the per-residue C_α_ displacement of PBD^PAK3^ between the aligned RhoJ and Cdc42 complexes. CAS was performed using BudeAlaScan (BAlaS). 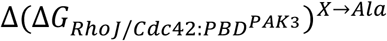was calculated using the following equation: 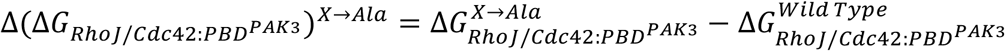 The brown dashed box highlights the 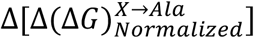and Cα displacements for PAK3 Leu72– Ser74. **(D)** CAS of RhoJ interface residues across triplicate 500 ns-long MD simulations. **(E)** Quantification of N-terminal contribution to RhoJ:PBD^PAK3^ interface. The N-terminal contribution to the RhoJ:PBD^PAK3^ interface was estimated by comparing full-length, 500ns MD-derived RhoJ:PBD^PAK3^ models against the corresponding models with a direct *in silico* truncation of the RhoJ N-terminal residues (Met1-Lys20). The N-terminus accounts for a buried surface area of 285.6 Å^2^ interface 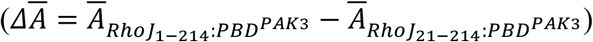 calculated by PDBePISA, representing ∼17.0% of total RhoJ:PBD^PAK3^ interface. **(F, G)** Structural and energetic impacts of RhoJ N-terminus on the RhoJ:PBD^PAK3^ interface within a physiological membrane context. Buried surface area **(F)** and predicted solvation free-energy gain upon interface formation (Δ^i^G) **(G)** were quantified across triplicate 500 ns on-membrane MD simulations using PDBePISA. N-terminal S-palmitoylation significantly reduced the buried interface area by an average of 213.4 Å^2^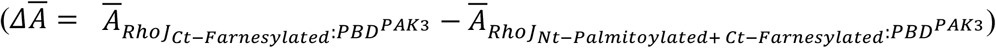(∼14.0% of the total RhoJ:PBD^PAK3^ interface area), and significantly decreased the magnitude of Δ^i^G by ∼17.7%, indicating less favourable interface formation. Data are presented as mean ± SEM (n = 3 independent simulations). Statistical significance was determined by a two-tailed Welch’s unpaired t-test (*p < 0.05).

### 3.2 N-terminal pocket formation with other RhoJ binding partners

To examine the contribution of the RhoJ N-terminal extension to effector binding more widely, we modelled the structures of RhoJ in complex with PAK1, N-WASP and BORG3. In our previous studies, N-WASP showed reduced binding to RhoJ compared to Cdc42 in endothelial cells, and BORG3 showed minimal binding compared to Cdc42 [16]. We included PAK1 as it has been reported to be a RhoJ binding partner in other cell types [38] and is highly homologous to PAK3 (84% identity).

Despite high sequence conservation in the core switch regions, the formation of the RhoJ N-terminal pocket was accompanied by significant structural divergence and a marked orientational shift in PAK positioning compared to Cdc42 complexes, most notably within the RhoJ:PBD^PAK3^ complex (Fig. 3A). Reproducible N-terminal wrapping and stable pocket formation occurred across all 3 MD-simulation replicates only for the RhoJ:PBD^PAK3^ complex. In contrast, while the representative RhoJ:PBD^PAK1^ structure, the one with the lowest-RMSD_Cα_, captures an apparent N-terminal pocket (Fig. 3B), this interface lacks structural stability, converging in only 1 of 3 replicates. Furthermore, the RhoJ N-terminus did not form the pocket structure with PBD^N-WASP^ and PBD^BORG3^, and did not stably contribute to the binding interface with the bound partner (Fig. 3C, D).

**Figure 3.**
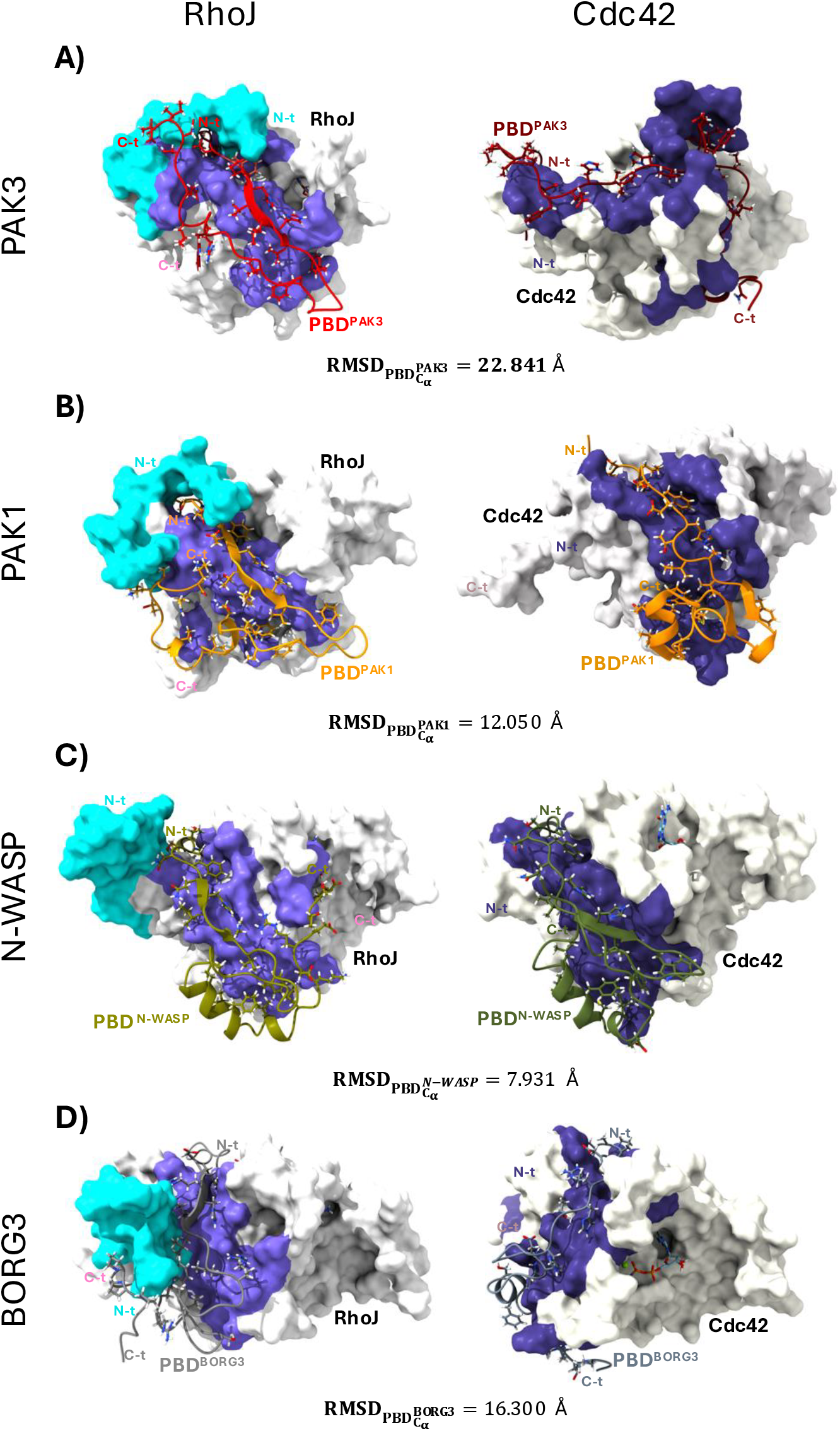
Comparison of the binding patterns of PBD between RhoJ and Cdc42. For all panels: Left, surface representation of the lowest-RMSD_Cα_ 500ns-long MD model of activated RhoJ (white; N-terminus Met1–Lys20, cyan; PBD-contact residues, medium slate blue). Right, reference structure of activated Cdc42 (ivory; PBD-contact residues, dark slate blue). Bottom, effector positional shifts quantified by RMSD_Cα_ when superimposing of RhoJ and Cdc42 complexes. **(A)** PBD^PAK3^ (red in RhoJ complex; dark red in Cdc42 NMR structure, PDB: 1EES). **(B)** PBD^PAK1^ (orange in RhoJ complex; dark orange in Cdc42 NMR structure, PDB: 1E0A). **(C)** PBD^N-WASP^ (olive in RhoJ complex; dark olive green in Cdc42 MD-model). **(D)** PBD^BORG3^ (gray in RhoJ complex; slate gray in Cdc42 MD-model). All models represent the activated (Mg^2+^-GppCp or GppNHp bound) state.

Notably, while PBD^PAK3^, PBD^PAK1^, PBD^N-WASP.^ shared similar binding surfaces on RhoJ, the specific orientation of PBD^PAK3^ differed substantially from its Cdc42-bound counterpart (Fig. 3, Fig. 2C). This structural displacement correlates with a unique energetic profile. BAlaS identified that residues E66–F84 of PAK3 comprise the primary energetic hotspots when bound to the RhoJ N-terminal pocket. In contrast, the Cdc42:PBD^PAK3^ interface is dominated by interactions within the D85–L102 region of PAK3 (Fig. 2C). Despite these unique interactions, RhoJ and Cdc42 complexes with PAK3 and PAK1 exhibited similar predicted solvation free-energy gains upon interface formation (Δ^i^G; Table 1). Their complexes with N-WASP also shared similar Δ^i^G values; however, RhoJ had a far lower |Δ^i^G| for BORG3 than Cdc42, in keeping with its lower apparent binding to this partner in endothelial cell lysates [16]. These data suggest that the RhoJ N-terminal pocket may not be the principal determinant of the overall interface energetics, because the absence of the interactions contributed by the N-terminal extension may be structurally compensated for by the G1 box and its adjacent residues present in the Cdc42:PBD^PAK3^ complex. This raises the possibility that the N-terminal pocket may not dictate binding affinity. Instead, it seems possible that the formation of RhoJ N-terminal pocket acts to selectively stabilize the formation of RhoJ:PAK3 complexes by decreasing the RhoJ:PAK3 dissociation rate. This unique allosteric site represents a novel target for selective pharmacological inhibition in endothelial cells, offering a strategy to disrupt tumour angiogenesis without cross-reacting with conserved, housekeeping Cdc42 pathways.

**Table 1.** Mean PDBePISA-predicted interface binding energies (Δ^i^G) for Cdc42 and RhoJ complexes with the indicated effector-binding domains (PBDs). Each value represents the mean Δ^i^G calculated from the 500-ns endpoint snapshot of each of three independent MD simulations per complex.

| | $\Delta^iG$ (kcal/mol) | |
| --- | --- | --- |
|  | Cdc42 | RhoJ |
| PAK3 | -27.7 | -20.0 |
| PAK1 | -22.5 | -20.5 |
| N-WASP | -18.6 | -20.6 |
| BORG3 | -20.3 | -7.1 |

### 3.3. Virtual screening identifies compounds fitting the RhoJ N-terminal pocket

To identify small-molecule inhibitors targeting the novel RhoJ N-terminal pocket, a structure-based virtual screen of a library of FDA-approved drugs was performed, using the equilibrated RhoJ-Mg^2+^-GppCP:PBD^PAK3^ complex model. To minimize the bias introduced by any single receptor conformation, each compound in the library was docked against all 5 representative RhoJ frames sampled across the equilibrated region of the MD trajectory at 200, 250, 300, 450 and 500 ns using BUDE [34], and predicted binding free energies were averaged across these 5 frames for each candidate. The global distribution of BUDE-predicted binding ΔG across the full drug library is displayed as a violin plot (Fig. 4, left), revealing a broad distribution of predicted scores. Compounds were subsequently ranked by mean predicted ΔG, and the top 20 candidates occupying the most favourable tail of the distribution are listed (Fig. 4, right).

**Figure 4.**
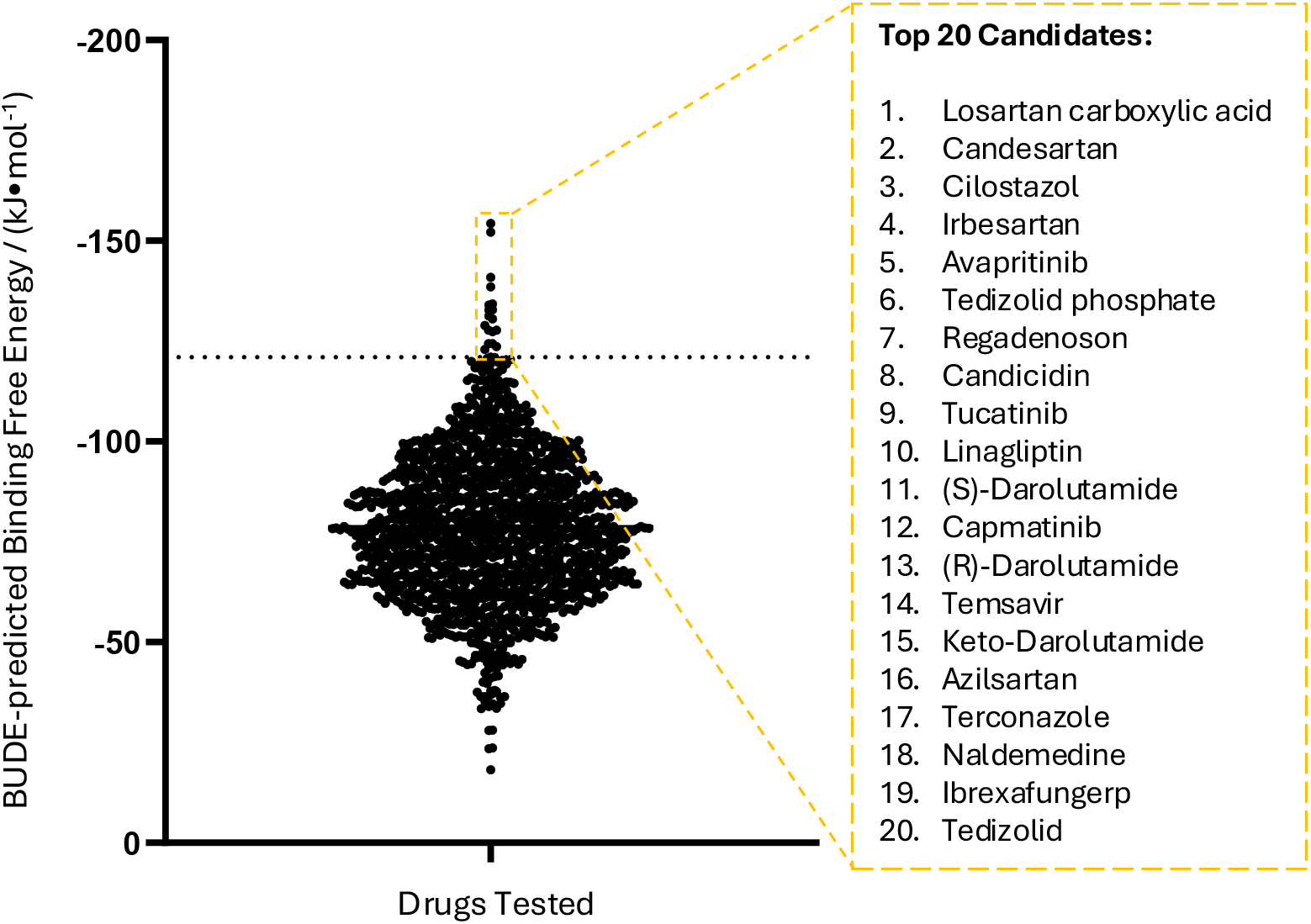
Structure-based virtual screening identifies potential inhibitors targeting the RhoJ N-terminal pocket. 5 equilibrated frames (200–500 ns) extracted from the activated RhoJ molecular dynamics (MD) trajectory (Fig. 1) were utilized for BUDE docking. BUDE-predicted binding free energies for each docked compound (ΔG, kJ·mol^−1^) were averaged across the 5 frames; data are presented as mean ± SEM. The violin plot illustrates the global binding energy distribution of the FDA-approved drug library, with the top 20 ranked candidates highlighted on the right.

### 3.4. Design of a cell-based BRET assay for RhoJ signalling

In order to validate the potential RhoJ inhibitors identified in the screen, we needed an assay for RhoJ effector binding. We constructed an enhanced bioluminescence resonance energy transfer (eBRET^2^)-based biosensor to monitor the RhoJ:PAK3 interaction. RhoJ was expressed as a fusion protein with the RLuc8 luciferase variant, and the PBD^PAK3^ was fused to the GFP variant GFP^2^. In the presence of the cell permeant luciferase substrate coelenterazine 400a, the light generated by the luciferase probe will excite the GFP^2^, but only if the two probes are in complex [39]. This assay has been used successfully as a cell-based screen for inhibitors of K-Ras [40] and has the advantage of allowing for screening in living cells.

To optimize resonance energy transfer efficiency while ensuring unconstrained formation of the RhoJ N-terminal pocket, biosensor configurations were designed based on the structural considerations of the RhoJ:PBD^PAK3^ interface. RLuc8 was inserted before the RhoJ C-terminal hypervariable region (HVR), and GFP^2^ was fused to either the N- or C-terminus of PBD^PAK3^, yielding two candidate orientations: Combination 1 (RhoJ-RLuc8-Ct : PBD^PAK3^-GFP^2^) and Combination 2 (RhoJ-RLuc8-Ct : GFP^2^-PBD^PAK3^) (Fig. 5A). Structural modelling by AlphaFold2-multimer indicated that both configurations provide sufficient conformational freedom for the RhoJ N-terminus to occupy the PBD^PAK3^ binding interface without steric hindrance from the fused fluorophores (data not shown). Empirical evaluation in HEK293T cells revealed that Combination 2 provided a significantly stronger BRET signal, superior dynamic range, and more robust performance with minimal variability compared to Combination 1. This was evidenced by a sharp decrease in the BRET ratio upon treatment with the Cdc42 inhibitor ML141 (Fig. 5B). Subsequent transfection optimization using a serial dilution of donor DNA (RhoJ_Q79L_-RLuc8-Ct) demonstrated that the BRET signal remained stable at a donor:acceptor (D:A) ratio of 1:10, while maintaining acceptable raw emission intensities (Fig. 5C).

**Figure 5.**
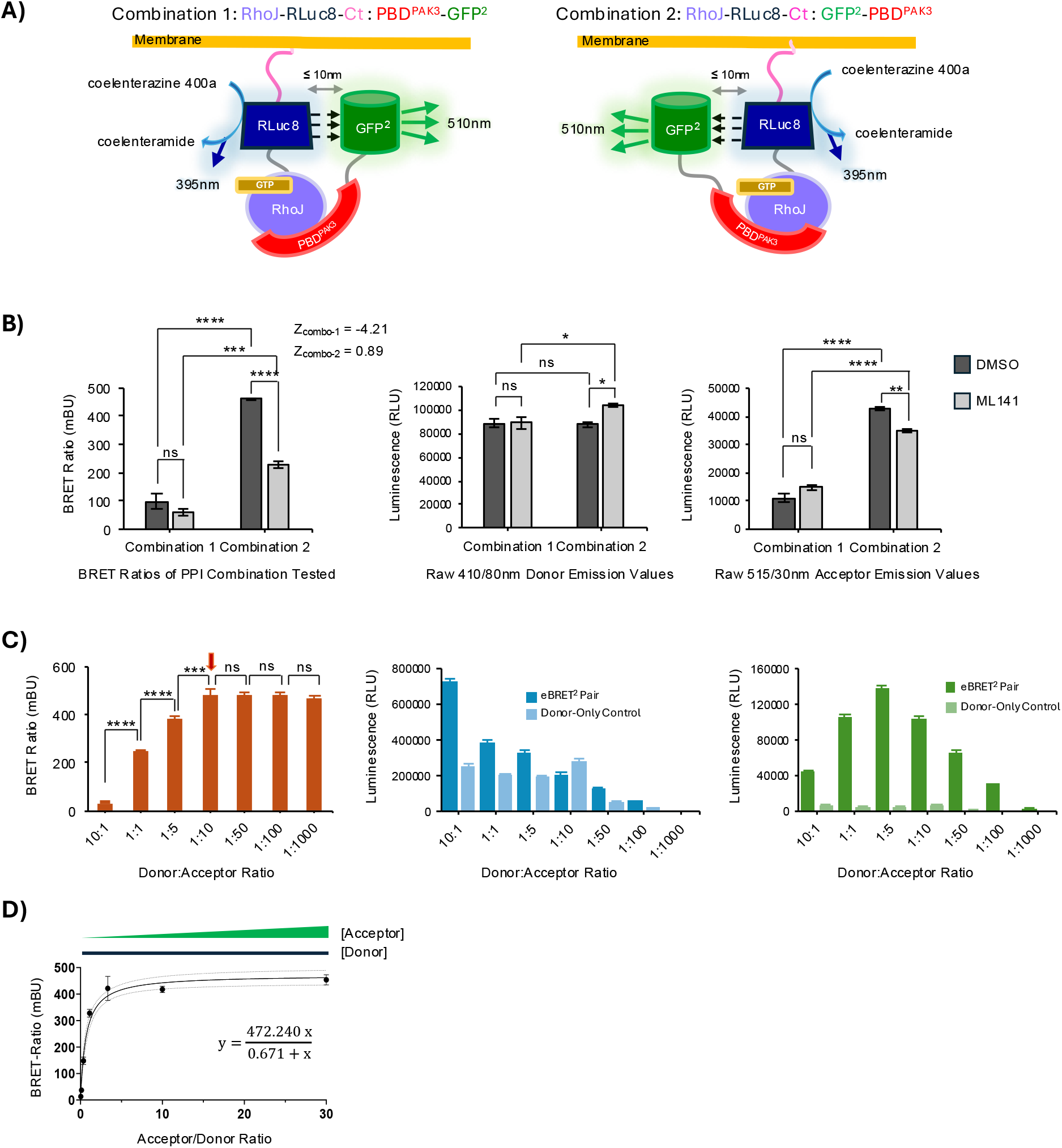
Construction and optimization of eBRET^2^-based RhoJ biosensors for screening. **(A)** Schematic representation of candidate eBRET^2^ biosensor configurations. Two distinct donor:acceptor orientations were designed to identify the optimal geometry for energy transfer: Combination 1 (RhoJ–RLuc8–Ct : PBD^PAK3^–GFP^2^) and Combination 2 (RhoJ– RLuc8–Ct : GFP^2^–PBD^PAK3^). Domain colors are assigned as follows: RhoJ (purple; C-terminal hypervariable region, hot pink); RLuc8 donor (blue); PBD^PAK3^ (red); GFP^2^ acceptor (green); and (GGGGS)_3_ flexible linkers (grey). **(B)** Orientation optimization of 2 possible eBRET^2^ pairs. HEK293T cells were co-transfected (1:10 donor:acceptor DNA ratio) and treated with DMSO or ML141. Corrected BRET ratios (mBU, left) and raw emissions for donor (410/80 nm, middle) and acceptor (515/30 nm, right) are shown. Combination 2 was selected for its superior dynamic range. **(C)** Transfection optimization by a serial dilution of donor DNA for the eBRET^2^ system. Combination 2 (RhoJ_Q79L_-RLuc8-Ct:GFP^2^-PBD^PAK3^) was titrated across indicated ratios while maintaining constant total DNA (3.2 μg) using empty pLVX-puro vector. Left, corrected BRET ratio (mBU); middle, raw donor emission (410/80 nm); right, raw acceptor emission (515/30 nm). **(D)** Corrected BRET saturation curve of the optimized RhoJ biosensor. HEK293T cells were transfected with fixed donor (RhoJ_Q79L_-RLuc8-Ct) and increasing amounts of acceptor (GFP^2^-PBD^PAK3^). Data were fitted to a nonlinear hyperbolic regression 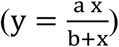, yielding a = BRET_max_ = 472.2 ± 18.1 mBU and b = BRET_50_ = 0.671 ± 0.116. Dotted lines indicate 95% confidence intervals. All data are presented as mean ± SEM. Statistics: one-way ANOVA with Bonferroni correction (C),and two-way ANOVA with Bonferroni correction (B) (ns = not significant; *p < 0.05; **p < 0.01; ***p < 0.001; ****p < 0.0001).

To confirm the specificity of the interaction, a BRET saturation assay was performed. The resulting saturable binding profile indicates that the signal is derived from a high-affinity protein-protein interaction rather than random collisions (Fig. 5D). The D:A ratio of 1:10 was further confirmed as the optimal ratio for inhibitor screening due to its low variance and high reproducibility. Collectively, these results establish the optimized RhoJ:PAK3 eBRET^2^ system (Combination 2) as a robust and sensitive tool for identifying small molecules targeting the RhoJ:PAK3 interaction.

### 3.5. The BRET assay confirms terconazole as a RhoJ inhibitor

The top 20 candidates were validated at 200 µM using the optimized RhoJ eBRET^2^ biosensor. Among these, terconazole emerged as the primary hit, demonstrating significantly higher inhibition of the RhoJ:PBD^PAK3^ interaction than the reference inhibitor ML141 (Fig. 6A). Dose-response characterization confirmed that terconazole inhibits the RhoJ biosensor signal in a dose-dependent manner with an apparent IC_50_ of 101 µM (Fig. 6B). Structural analysis of the predicted binding mode revealed that terconazole occupies the N-terminal allosteric pocket, effectively sequestering the extension and preventing effector recruitment (Fig. 6C). Unlike previously reported inhibitors that target the G-domain interface, terconazole’s binding is stabilized by a network of hydrophobic contacts within the RhoJ-specific N-terminal region (Fig. 6D). With the exception of RhoJ Asp44, these interactions overlap extensively with the residues comprising the natural PBD^PAK3^ interface (data not shown), suggesting that terconazole inhibits RhoJ activity by competing with PAK3 for access to the N-terminal pocket. Finally, selectivity for RhoJ was assessed against Cdc42, using the well-characterised G-LISA assay. Purified constitutively active Cdc42 was treated with terconazole (10 or 100 µM) for 15 min, and the binding of the Cdc42 to immobilised PBD^PAK1^ was measured by ELISA. Terconazole had no effect on Cdc42 binding to PBD^PAK1^ at either concentration relative to the vehicle control (Fig. 6E), supporting selectivity for RhoJ over Cdc42 and consistent with recognition of structural features unique to the RhoJ N-terminal pocket. Collectively, these findings provide a structural basis for terconazole inhibitory activity and identifies terconazole as a lead scaffold for targeting this novel allosteric site in RhoJ.

**Figure 6.**
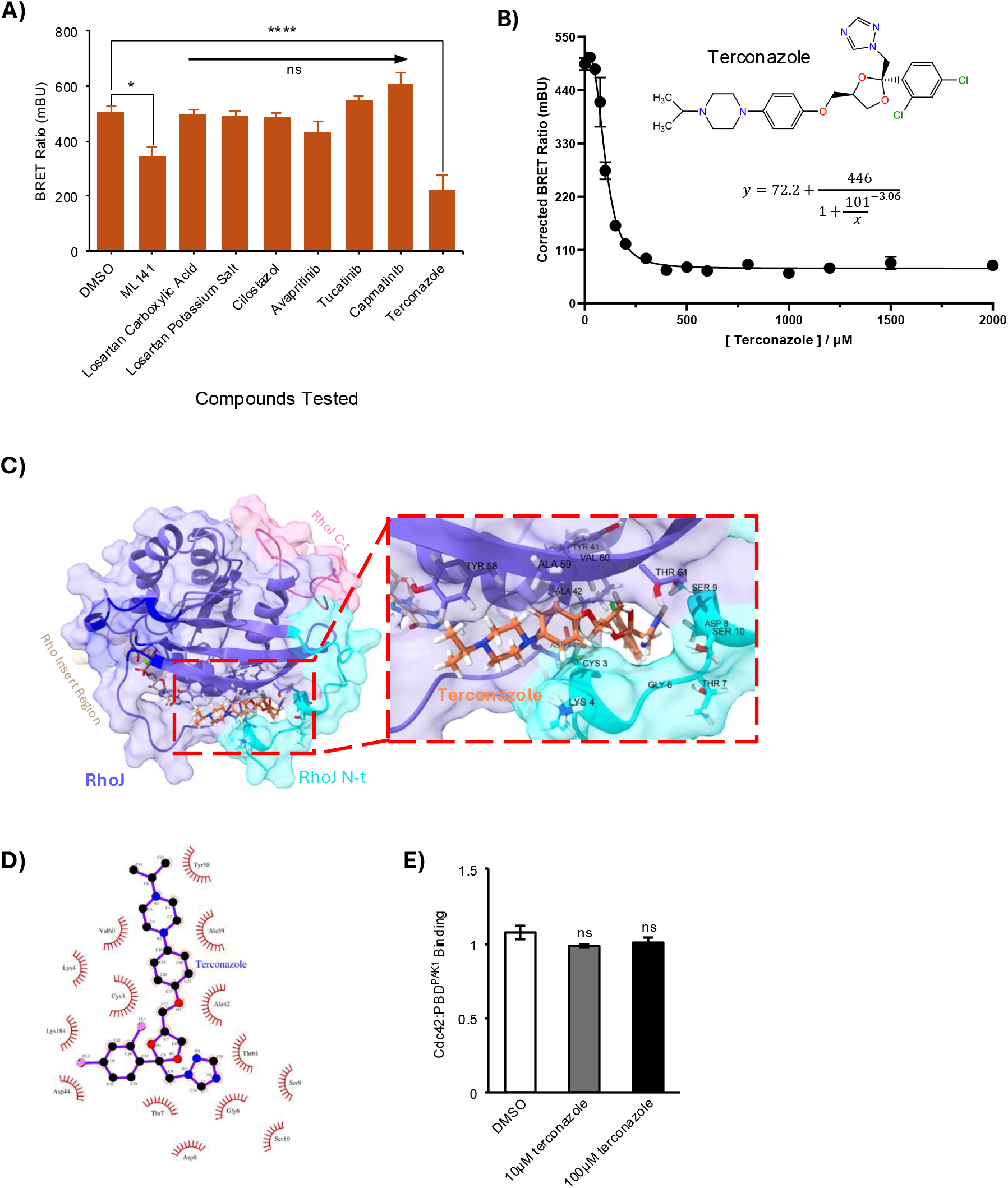
Experimental validation and structural characterization of terconazole as a selective RhoJ inhibitor. **(A)** eBRET^2^-based screening of top virtual hits targeting RhoJ N-t pocket. HEK293T cells expressing the RhoJ biosensor (RhoJ_Q79L_–RLuc8–Ct : GFP^2^–PBD^PAK3^, donor:acceptor ratio = 1:10) were treated with candidates (200 µM), DMSO (1% v/v; vehicle control), or ML141 (20 µM; inhibition control) for 30 min. Statistics: one-way ANOVA with Bonferroni post-hoc test (ns, not significant; *p < 0.05, ****p < 0.0001). n = 3 independent experiments. **(B)** Dose-response characterization of terconazole. Corrected BRET ratios were fitted to a 4-parameter logistic regression, 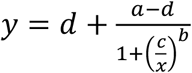, yielding a = BRET_max_ = 518 ± 12.9 mBU, b = ™3.06 ± 0.301, c = IC_50_ = 101 ± 6.80 μM, and d = 72.1 ± 2.27 mBU. Dotted lines indicate 95% confidence intervals of the fitted curve. n = 3 independent experiments. **(C)** Predicted binding mode of terconazole within the RhoJ N-terminal pocket. Terconazole (coral sticks) was docked into a representative MD snapshot (450 ns, with a BUDE-predicted ΔG closest to the mean across the 5 dockings). RhoJ is shown in surface representation (medium slate blue; N-terminus, cyan; Rho-insert region, tan; C-terminus, hot pink; previously reported allosteric pocket residues, blue). Mg^2+^ and GppCp are shown as lime sphere and tan sticks, respectively. Inset: Detailed view of the binding site with labelled contacting residues. **(D)** 2D interaction fingerprint of the terconazole docking pose from (C), illustrating hydrophobic contacts (red radial sectors) with surrounding residues. **(E)** Selectivity assessment of terconazole against Cdc42. Purified constitutively active Cdc42 was incubated with vehicle [5% (v/v) DMSO] or terconazole for 15 min. Binding of Cdc42 to immobilised PBD^PAK1^ was quantified by G-LISA assay. Terconazole produced no significant alteration in Cdc42 binding to PBD^PAK1^ compared to vehicle control. n = 4 independent experiments. Statistical significance was assessed using a Geisser-Greenhouse-corrected one-way repeated-measures ANOVA followed by Dunnett’s post-hoc test against the vehicle control (ns, not significant). Data are mean ± SEM.

## 4. Discussion

This study identifies a previously unrecognised mechanism by which the RhoJ N-terminal extension contributes to effector engagement and exploits this feature as a route to selective pharmacological targeting. RhoJ is closely related to Cdc42 across the conserved GTPase core, which has made selective inhibitor design challenging. Our findings indicate that this difficulty may be overcome by targeting a structural element that is absent from Cdc42 – the RhoJ-specific N-terminus. In the context of PAK3 binding, this region formed a defined pocket at the RhoJ interface. This pocket made a substantial contribution to interface area and predicted binding energy, suggesting that the N-terminal extension is an active determinant of RhoJ effector recognition.

A notable feature of the modelling is that pocket formation was most stable in the RhoJ:PAK3 complex. Although RhoJ can engage several Cdc42-family effectors, including PAK1 and N-WASP, the simulations suggest that these interactions are not structurally equivalent. The RhoJ N-terminus consistently wrapped across the PAK3 interface, whereas comparable behaviour was not reproduced with N-WASP or BORG3 and was less stable with PAK1 (Fig. 3). These findings support a model in which the RhoJ N-terminus contributes to effector selectivity by altering the geometry and stability of the bound complex, rather than simply increasing global affinity. This may be particularly relevant in endothelial cells, where PAK3 appears to represent a major RhoJ signalling partner [16]. By shaping the orientation or lifetime of RhoJ complexes, the N-terminal region could provide a mechanism through which RhoJ generates signalling outputs distinct from those of Cdc42, despite strong conservation of the switch-region binding surface.

Although terconazole demonstrated superior inhibition compared to ML141, we note that at 101 µM, it is relatively weak binder. This, and the lack of specific hydrogen bonds between terconazole and the RhoJ N-terminus in our docking models (Fig. 6D), suggest that binding is primarily driven by hydrophobic contacts. While this may imply a lack of the strict spatial constraints typically afforded by polar networks, a hydrophobic-driven binding mechanism offers distinct thermodynamic advantages, notably a minimal entropic penalty achieved via the displacement of ordered water molecules from the allosteric pocket upon ligand engagement. Additionally, such hydrophobic character is highly favourable for cellular distribution and membrane permeability, which are critical parameters for targeting intracellular small GTPases. Further, the N-terminal pocket represents a distinct, targetable site for the development of selective anti-angiogenic agents that bypass the challenges of G-domain conservation, offering a potential path toward more potent and selective RhoJ inhibitors for anti-angiogenic therapies. Therefore, while terconazole itself may lack the potency required for immediate clinical translation, achieving competitive inhibition at this unique allosteric site serves as a critical proof-of-concept. Its unique hydrophobic scaffolding provides an optimized starting point for structure-based hit optimization aimed at enhancing binding affinity while maintaining a low desolvation penalty. Future strategies could focus on modifying the triazole scaffold that satisfy the directional hydrogen-bonding requirements of the allosteric pocket residues to not only optimize the physicochemical profile by improving aqueous solubility, which is typically limited for this hydrophobic scaffold, but would also simultaneously provide a favourable enthalpic driving force to enhance target potency.

Taken together, our findings define the RhoJ N-terminal pocket as a selective allosteric site and demonstrate that it can be targeted by small molecules. More broadly, they suggest that lineage-specific extensions in closely related small GTPases may create exploitable surfaces for selective drug discovery. For RhoJ, this provides a rational path towards a new class of anti-angiogenic inhibitors acting downstream of multiple pro-angiogenic signals.

## CRediT authorship contribution statement

Wenxuan Chen: Conceptualization (supporting); investigation (lead); data curation (lead); methodology (equal); formal analysis (lead); visualization (lead); writing—review and editing (equal); writing—original draft (equal).

Deborah Karen Shoemark: Supervision (supporting); methodology (supporting); formal analysis (supporting).

Robin A Corey: Formal analysis (supporting); writing—review and editing (equal); writing— original draft (equal).

Tatyana N. Andrienko: Investigation (supporting); methodology (supporting).

Harry Mellor: Conceptualization (lead); funding acquisition (lead); methodology (equal); supervision (lead); writing—original draft (equal); writing—review and editing (equal); project administration (lead).

## Declaration of completing interest

The authors declare that they have no known competing financial interests or personal relationships that could have appeared to influence the work reported in this paper.

## Acknowledgements

W. Chen was supported by a PhD scholarship from the Chinese Scholarship Council.

## Data availability

All the data supporting the findings of this study are included in the paper. The computational structural models associated with this study are available through the Open Science Framework at https://doi.org/10.17605/OSF.IO/MYDEU.

